# Biomaterial based implants caused remote liver fatty deposition through activated blood-derived Kupffer cells

**DOI:** 10.1101/2023.04.12.536663

**Authors:** Zhi Peng, Chang Xie, Shucheng Jin, Jiajie Hu, Xudong Yao, Jinchun Ye, Xianzhu Zhang, Jia Xuan Lim, Bingbing Wu, Haoyu Wu, Renjie Liang, Ya Wen, Jiahui Huang, Xiaohui Zou, Hongwei Ouyang

## Abstract

Understanding the foreign-body response (FBR) of biomaterials is a prerequisite for the prediction of its clinical application, and the present assessments mainly rely on in vitro cell culture and in situ histopathology. However, remote organs responses after biomaterials implantation is unclear. Here, by leveraging body-wide-transcriptomics data, we performed in-depth systems analysis of biomaterials - remote organs crosstalk after abdominal implantation of polypropylene and silk fibroin using a rodent model, demonstrating local implantation caused remote organs responses dominated by acute-phase responses, immune system responses and lipid metabolism disorders. Of note, liver function was specially disturbed, defined as hepatic lipid deposition. Combining flow cytometry analyses and liver monocyte recruitment inhibition experiments, we proved that blood derived monocyte-derived Kupffer cells in the liver underlying the mechanism of abnormal lipid deposition induced by local biomaterials implantation. Moreover, from the perspective of temporality, the remote organs responses and liver lipid deposition of silk fibroin group faded away with biomaterial degradation and restored to normal at end, which highlighted its superiority of degradability. These findings were further indirectly evidenced by human blood biochemical examination from 141 clinical cases of hernia repair using silk fibroin mesh and polypropylene mesh. In conclusion, this study provided knowledge of biomaterials-body interactions. It is of great important for future development of biomaterial devices for clinical application.

**One Sentence Summary:** Abdominal local biomaterials implantation induces remote organ fatty deposition through activated blood-derived Kupffer cells.

## INTRODUCTION

Biomaterials play an important role in tissue engineering and clinical biomedical devices*(1)*. A key concept in selecting proper biomaterial is to induce no or minimal foreign-body response (FBR) from the recipient*(2)*. However, all biomaterials implantation will inevitably initiate the host FBR*(3)*. This response to biomaterials can affect the performance of the implants during application, and even lead to a range of complications such as infection, chronic inflammation, tissue adhesion and chronic pain*(4, 5)*. Therefore, assessing the host FBR is still the most important field of research in biomaterials design and application.

There has been a huge amount work in the field of developing methods to assess the host FBR of biomaterials*(6)*, and many different tests have been performed as recommended by various regulatory agencies*(7)*. A lot of studies have been conducted in vitro through the evaluation of cell viability, morphology, adhesion, proliferation and migration*(8, 9)*. However, they failed to accurately model the dynamic and extremely complex environment because cells and body fluids never stay static in vivo*(7)*. In vivo, most previous studies have focused on implanted site with the use of histological assessment*(10–12)*. Therefore, adverse local complications such as fibrotic capsule*(13, 14)* and inflammatory response*(15–17)* have been widely described, while systemic evaluating the host FBR from remote organs responses perspective remains an emerging field.

In general, tissue engineering associated biomaterials broadly classified by biological or synthetic*(11)*. Non-degradable synthetic biomaterials, such as polypropylene (PP), provide strong reinforcement for soft tissues repair and preserve stable quality in the long-term*(18)*. Additionally, the PP mesh has been used for more than 50 years in abdominal hernia repair in the form of a patch sutured to the abdominal wall*(19)*. In contrast, biological scaffolds, such as natural derived silk fibroin (SF), have been intensively explored as biomaterials for tissue engineering in recent years benefit from its excellent mechanical and biological properties*(20)*. Presently, SF-based biomaterials have been widely applied in several tissue regenerations including bone*(21)*, cartilage*(22)*, vascular*(23)*, skin*(20)* and hernia*(6, 24)*.

However, whether they are natural or synthetic, a series of articles has been reported that locally implanted biomaterial devices triggered systemic responses. After the subcutaneously implantation of gelatin-based biomaterial on the mice back, activation of NF-kB was observed in remote major organs including the lung, spleen and small intestine*(25)*. After volumetric muscle loss (VML) injury followed by submucosal-intestine ECM (SIS-ECM) implantation on mice, mature B cells and antigen presentation were observed in the draining inguinal lymph nodes and spleen*(26)*. Autoinflammatory/autoimmunity syndrome (ASIA; Shoenfeld’s syndrome) represents a systemic illness with complications such as cognitive impairment, joint pain, myalgias, fever, fatigue, dry eyes and mouth in patients*(27)*. Tervaert et, al. reviewed ASIA induced by biomaterials with clinical case reports including silicone breast implants*(27)* and PP meshes implantation (pelvic organ prolapse surgery and hernia repair)*(28)*. Our previous research demonstrated that after PP meshes abdominal implantation in mice, both the innate and adaptive system in blood were activated*(6)*, but the specific effect of local biomaterials implantation on remote major tissues and organs in the whole-organism scale still remain unknown.

In the present work, we especially focused on the remote organs responses after biomaterials implantation in vivo. Synthetic PP and biological SF, were typically selected as model biomaterials in abdominal implantation, because of their long history of hernia repair’s clinical use and commercial potentiality. Based on the disturbed transcriptome expression pattern of remote major organs induced by biomaterials implantation, we observed liver lipid deposition and validated local-blood-organ mechanisms mediated by circulating monocyte pool. Moreover, we found that the remote organs responses and liver lipid deposition were attenuated in SF group after a long-term duration. These findings were further indirectly verified by the blood biochemical liver function indicators (ALT, AST) in a clinical trial with PP mesh or SF mesh for hernia repair. Concepts learned from these results might be referenced by clinical applications when choosing the type of implants for patients and illuminate general guidelines by which to assess whole-organism host-biomaterials interactions.

## RESULTS

### Subhead 1: Whole-organism gene expression analysis revealed remote organs responses by biomaterials implantation

To mimic the routes of clinical usage, PP and SF meshes were implanted into the abdominal wall of Sprague-Dawley rats and sham operated rats as control group, followed by investigating the host gene expression dynamics across the organism after implantation. We performed whole-organism-wide RNA sequencing (RNA-seq) on 7 major organs/tissues (blood, brain, heart, kidney, liver, lung, spleen) collected at 3 different time points (2w, 6w and 24w) based on different host response phases (Fig. 1A). In all tissues, gene expression changed after two types of mesh implantation (Fig. 1). On average, 49 differentially expressed genes (DEGs) in PP group and 55 DEGs in SF group after implantation compared to tissue - matched controls. The amplitude of the host response varied among tissues and time points, and blood at 2w showed the highest DEGs in both PP (388) and SF (350) groups (Figure 1B; Supplementary Data 1 and 2). In SF group, blood and liver displayed the highest number of DEGs. The liver had the second lowest number of DEGs after PP implantation (Liver 8, Supplementary Data 1). However, when the highest fold change genes were averaged across all tissues, the median of the liver was high (Fig. 1C, top; Supplementary Data 3 and 4), considering that the liver was a major organ disturbed by biomaterials implantation according to DEGs’ numbers and fold changes. Analysis of the hallmark functional gene sets revealed that the implantation altered more than one hundred pathways (Supplementary Data 5 and 6). Even though a few of differentially expressed genes exists, alterations in the pathways were still significant. For instance, brain at 2w in SF group only exhibited 5 differential genes with FDR <0.1, but significantly altered 21 hallmark functional gene sets.

**Fig. 1.**
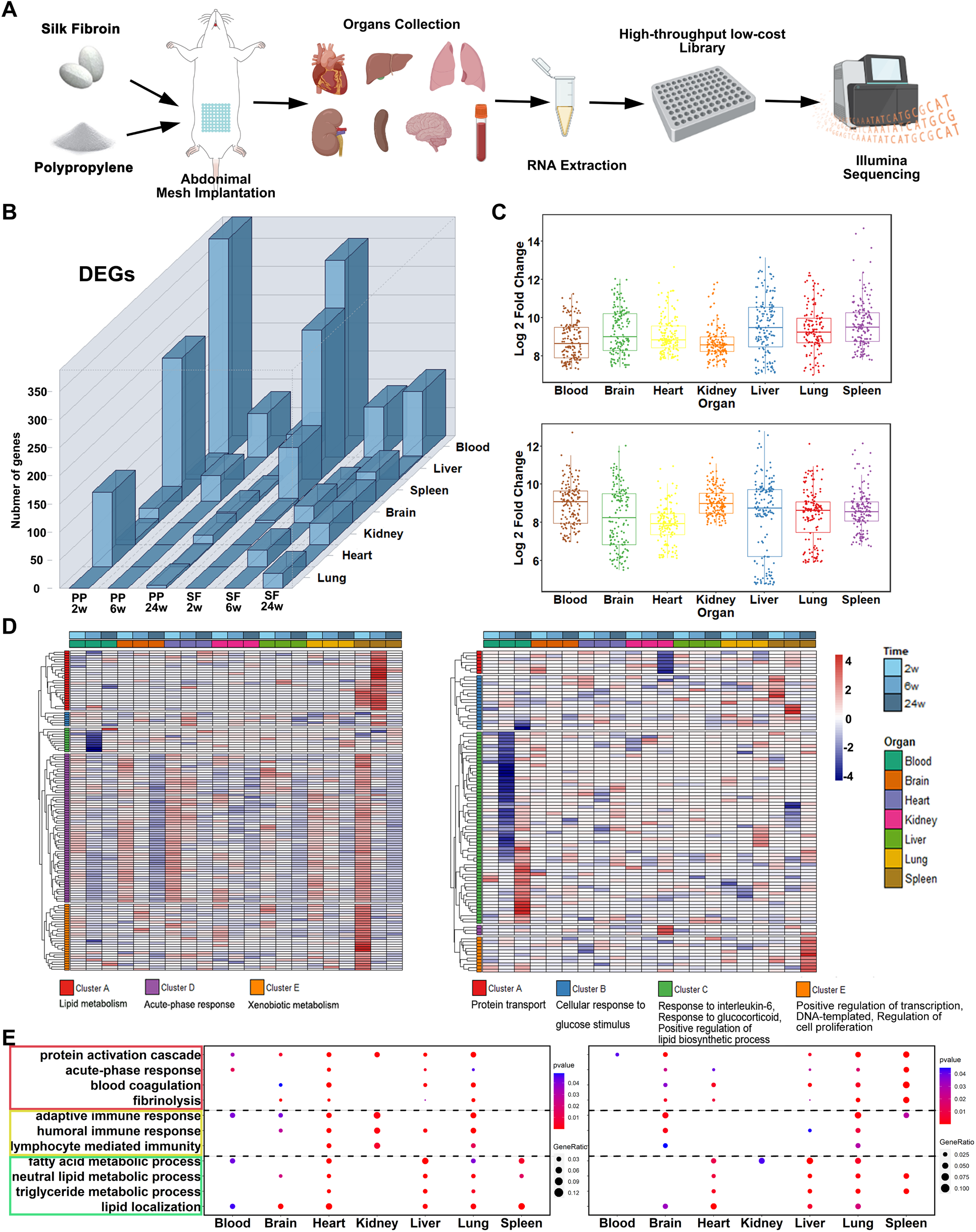
Whole-organism gene expression analysis reveals biomaterials-induced systemic response. **A**, Schematic illustration of the study: organs/tissues were collected after Control/PP/SF treatment, and used for RNA extraction, library preparation, sequencing, and downstream bioinformatic analysis. **B**, Number of differentially expressed genes (FDR < 0.1) in organs/tissues after PP and SF implantation. **C**, Log2 fold changes of the most increased gene sets (by p value) after PP (left) and SF (right) implantation. For each organs/tissues, the top 50 gene sets were included. **D**, Heatmap showing log2 fold changes for the top differentially expressed unique genes (FDR < 0.01 in at least one condition) after PP (left) and SF (right) implantation. **E**, GO enrichment analysis of genes that are expressed with a fold change > 2 in at least two time points. PP (left) and SF (right).

To investigate the most significant implantation-related transcripts changes in tissues, we summarized the fold change of DEGs (with FDR<0.01) across the entire samples (Fig. 1D). By clustering genes, we obtained five gene sets in each group that showed correlated expression (Supplementary Data7 and 8). In PP group, a large group (cluster A, D and E) differentially increased in spleen after implantation. These genes clustered in the lipid metabolism, acute-phase response and xenobiotic metabolism (Fig. 1D, left). In SF group, a large group (cluster C) differentially expressed in blood after implantation, which clustered in the inflammatory response (response to interleukin-6 and glucocorticoid). This cluster also participated in the positive regulation of lipid biosynthetic process, hinting that there was a lipid disorder in the plasma. Other notable groups included genes implicated in protein transport (cluster A) in the kidney, cellular response to glucose stimulus (cluster B) and positive regulation of transcription, DNA-templated and regulation of cell proliferation (cluster E) in the spleen. In addition, we noticed that some genes only altered at multiple time points in one organ (Fig. 1D). To test the similarity of differentially expressed gene within one single organ or multi-organs, we pairwise compared genes with fold change greater than 2 (Supplementary Fig. 1). In PP group, the lung, liver and heart showed a similar response pattern to implantation. However, SF group showed a high similarity within organs.

To determine the functional categories of higher expression gene in multi-organs during post-implantation, Gene Ontology (GO) enrichment analysis were performed in genes with fold change greater than 2 in at least two time point (corresponds to the overlap of Venn-diagrams in Supplementary Fig. 2). Some GO terms were extensively enriched in several organs in both groups (Fig. 1E). For example, ‘‘Acute-phase response” (red rectangle) were enriched in several organs such as the brain, heart, liver and lung. “Immune response” (yellow rectangle) related GO terms rose mainly in the brain and lung. Of note, several members of the lipid transport and metabolic process (green rectangle) were expressed at higher levels in the heart, liver and lung (Fig. 1E), suggesting lipid homeostasis was affected extensively. Together, extensive genes were altered across all remote organs/tissues, including immune response, acute-phase response and lipid metabolism, after local implantation.

### Subhead 2: Local biomaterials implantation raised liver lipid droplets deposition

To determine the effect of biomaterials implantation on the protein levels, we next performed whole remote organs histological evaluation with no abnormalities found (Supplementary Fig. 3 and 4A). Blood samples were collected and analyzed by blood routine and chemistry examination following mesh implantation (Fig. 2A). Mean corpuscular hemoglobin concentration (MCHC) was decreased in both group after 2w implantation. Platelet (PLT) was increased after SF (2w) and PP (24w) implantation. Of note, liver function index (alkaline phosphatase, ALP, alanine aminotransferase, ALT, triglyceride, TG and total protein, TP) were disturbed extensively, indicating that liver function may be impaired by mesh implantation. In addition, the hepatic lesion of the two groups showed time differences. In 2w and 6w, liver function index was disturbed in both PP (ALP, ALT and TG (p = 0.0628)) and SF (ALT) group. However, only PP (TP and ALT (p = 0.0524)) group was disturbed at 24w (Fig. 2A), proposing that hepatic lesion may attenuated in SF after 24w implantation.

**Fig. 2.**
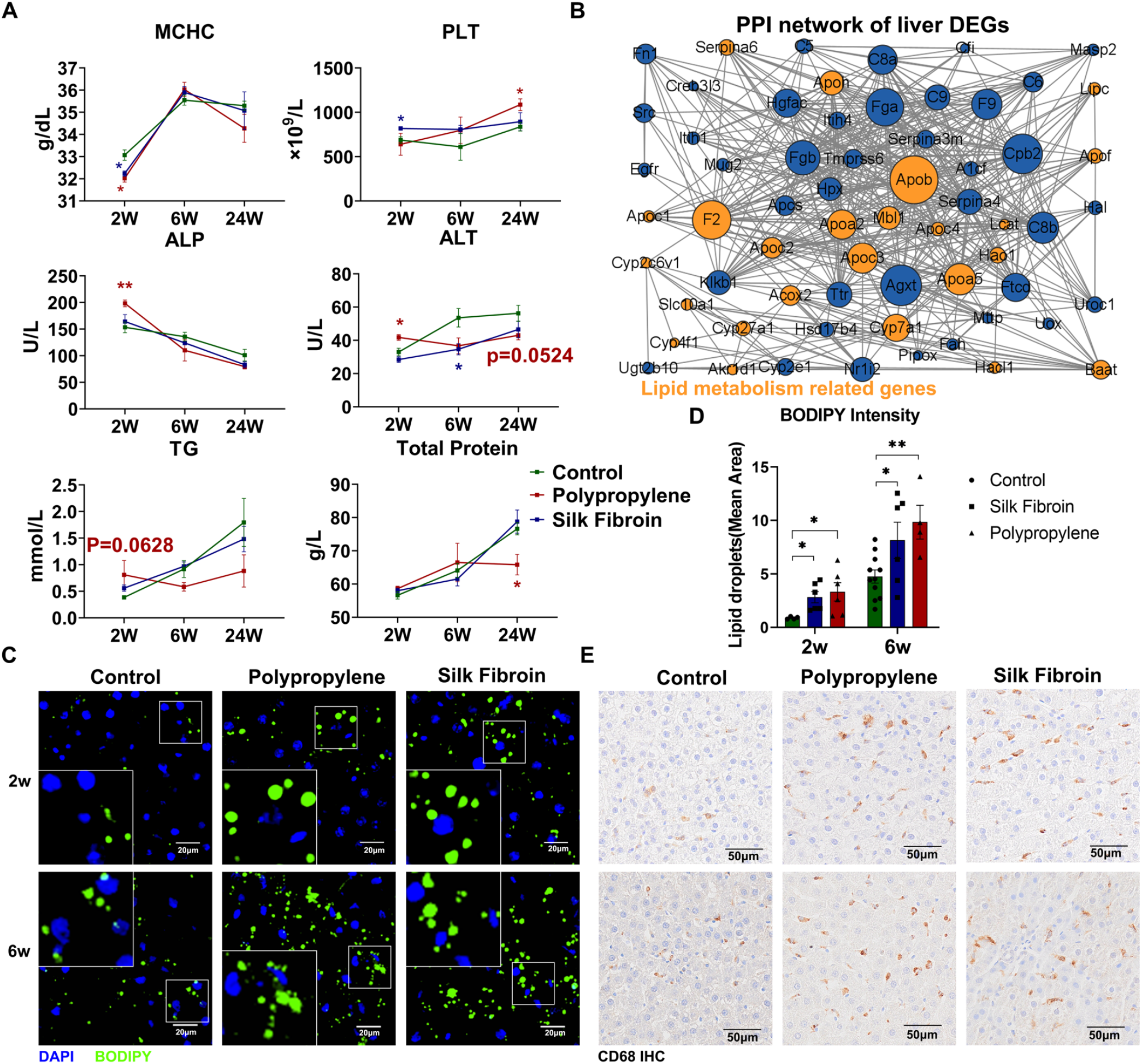
Local biomaterials implantation induced liver lipid droplets deposition. **A**, Blood routine and serum chemistry analysis of rats treated with PP and SF implantation. Data are mean ± SEM (n = 4). *p < 0.05, **p < 0.005, Student’s t test. **B**, StringDB protein-protein association network for all differentially expressed genes in liver after implantation (orange, lipid metabolism process related genes). **C, D**, Fluorescence staining of the liver lipid after implantation. Data are mean ± SEM (n = 4-11). *p < 0.05, Student’s t test. **E**, Immunohistochemistry staining showing Kupffer cell marker CD68 in liver sections.

Liver plays as a center pleiotropic endocrine organ affecting organism metabolism*(29)* and major changes occur at the transcriptome level (Fig. 1 and Supplementary Fig. 4B). So, we constructed a protein-protein interaction network analysis using the intersection of all differentially expressed genes in the liver after implantation, and the results showed that 38.7% of these genes belonged to the lipid metabolism related process (orange circles) (Fig. 2B). Next, in vivo validation of the mesh implantation induced lipid metabolism disturbance in the liver were conducted. The fluorescence and Oil red O staining in the liver demonstrated significant increased liver lipid droplet deposition in both PP and SF groups compared to the control group in 2w and 6w (Fig. 2C, D and Supplementary Fig. 4C). To explore the underlying mechanism of biomaterials induced liver lipid deposition, we attempted to assess the liver inflammation level through hepatic macrophages, specifically Kupffer cells (KCs)*(30)*. The results revealed that the fluctuation of KCs was highly consistent with lipid deposition in Fig. 2C (Fig. 2E), indicating that the up-regulated inflammation from KCs may be responsible for lipid accumulation. Taken together, these data verified that local biomaterials implantation can cause remote liver lipid droplets deposition.

### Subhead 3: Hepatic monocyte-derived KCs (moKCs) mediated lipid droplets deposition against biomaterials implantation

During the process of metabolic-associated fatty liver disease (MAFLD), resident Kupffer cells (KCs) are lost with progression and replaced by monocyte-derived KCs (moKCs)*(30)*. moKCs in a pro-inflammatory state contributed to the progression of non-alcoholic steatohepatitis (NASH)*(31)*. So, flow cytometry analyses were conducted to analyze the KCs composition of the liver. Gating strategy was used to identify CD45^high^ Ly6G^low^ CD64^high^ F4/80^high^ CD11b^low^ resident KCs and moKCs (Fig. 3A(a) and Supplementary Fig. 5A) as previous studies*(30–32)*.

**Fig. 3.**
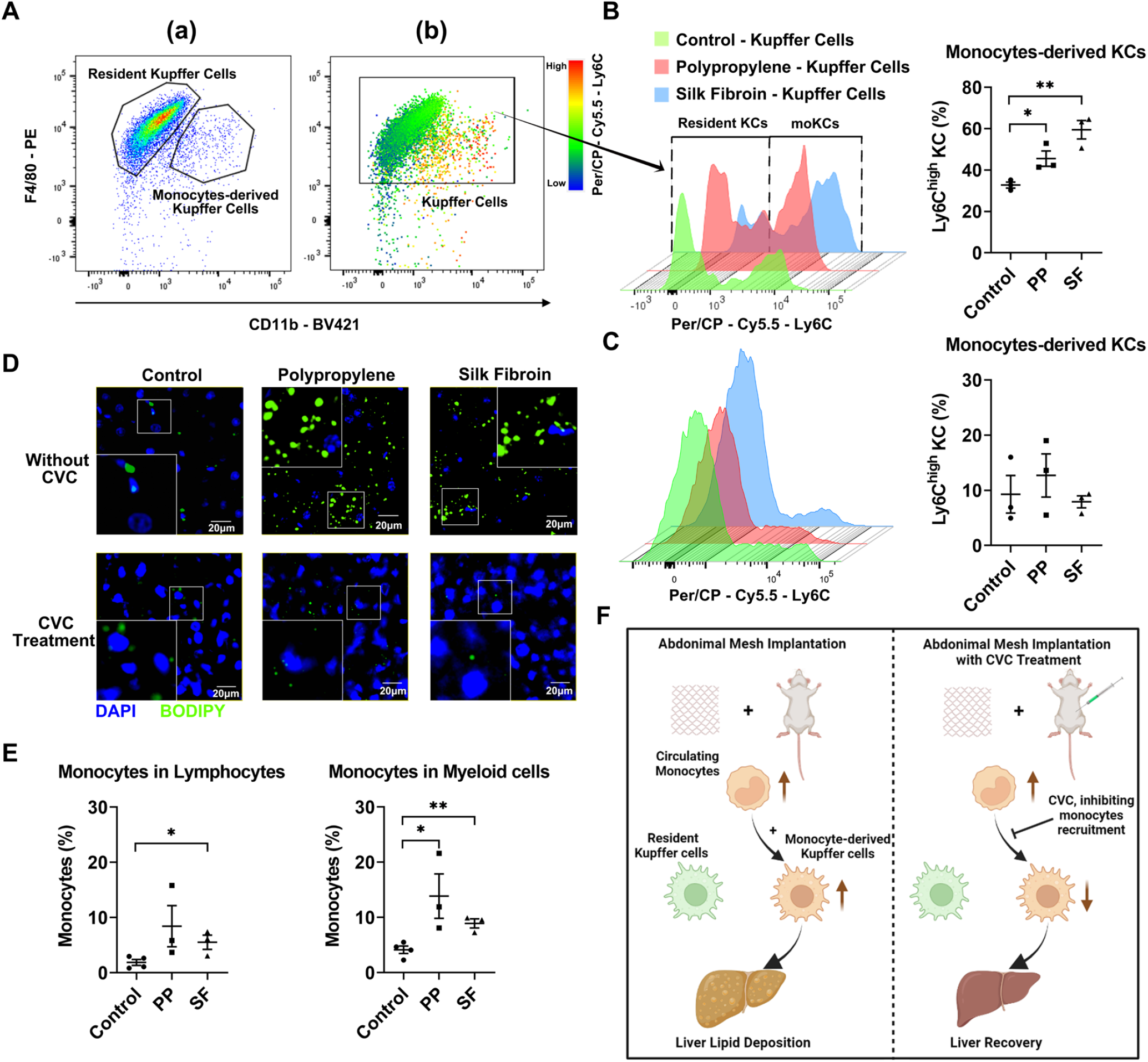
moKCs mediated biomaterials induced lipid droplets deposition. **A**, Gating strategy used to identify the KCs type in the liver (a) and the heatmap of Ly6C expression level (b). **B**, Flow cytometry analysis (left) and quantification (right) of liver moKCs and resident KCs after biomaterials implantation. Data are mean ± SEM (n = 3). *p < 0.05, **p < 0.005, Student’s t test. **C**, Flow cytometry analysis (left) and quantification (right) of liver moKCs after biomaterials implantation with CVC treatment. Data are mean ± SEM (n = 3). **D**, Fluorescence staining of the lipid droplet in liver sections after biomaterials implantation with/without CVC treatment. **E**, Flow cytometry analysis of blood circulating monocytes proportion in the lymphocytes (left) and myeloid cells (right). Data are mean ± SEM (n = 3-4). *p < 0.05, **p < 0.005, Student’s t test. **F**, Schematic illustration of the moKCs mediated liver lipid droplets deposition.

Cell surface markers Ly6C is high expressed in circulating monocytes and the heatmap of the Ly6C expression level indicated that Ly6C was capable of discriminating resident KCs and moKCs (Fig. 3A(b)). So, in this study, we identified resident KCs as Ly6C ^low^ and moKCs as Ly6C^high^. With the use of flow cytometry analyses, results presented that the proportion of moKCs increased after PP and SF implantation in liver (Fig. 3B), indicating that moKCs may mediate local biomaterials implantation induced liver lipid deposition. Liver moKCs are recruited through the chemokine receptor C-C motif chemokine receptor 2 (CCR2). So, cenicriviroc (CVC), a chemokine receptor CCR2 antagonist, was used intraperitoneally aiming to therapeutically inhibit liver monocyte recruitment*(33)*. The results demonstrated that CVC treatment specifically blocked up-regulated liver moKCs after biomaterials implantation (Fig. 3C). Most importantly, CVC treatment rescued the liver lipid deposition from biomaterials implantation (Fig. 3D). Besides, flow cytometry analyses of peripheral blood mononuclear cells (PBMC) confirmed that the proportion of circulating Ly6C^hi^ monocyte was significantly increased in both lymphocytes (CD45^+^) and myeloid cells (CD45^+^CD11b^+^) population (Fig. 3E), which were consistent with the CytoF results in a previous study*(6)*. These results confirmed that the increased moKCs were derived from up-regulated blood circulating Ly6C^hi^ monocytes. In conclusion, blood circulating monocytes derived liver moKCs mediated local biomaterials induced liver lipid droplets deposition (Fig. 3F).

### Subhead 4: Biomaterials-induced immune changes promoted blood circulating monocytes

Next, we sought to investigate how the local biomaterials implantation led to an increase in blood circulating monocytes. At first, the cytokines and chemokines in the blood serum were screened by luminex assay. Cxcl16, Ccl12, Cxcl5 were up-regulated in two experiment groups compared with control group (Fig. 4A, B and Supplementary Fig.6A). Ccr2 is a chemotactic receptor that mediates monocyte bone marrow egress, and Ccl12 binds to Ccr2, therefore Ccl12 may be a key molecule mediating the elevation of blood monocytes. Then, to investigate the source of Ccl12 protein, we also detected the cytokines and chemokines in the implanted site muscle. 19 out of 22 cytokines and chemokines were up-regulated in two experiment groups compared with control group (Fig. 4C and Supplementary Fig. 6B). Specifically, the protein levels of Ccl12 in the implanted site were much higher (> 10 fold) than in the blood serum (Fig. 4B, D), suggesting that they resourced from the implanted site and released into the blood circulation.

**Fig. 4.**
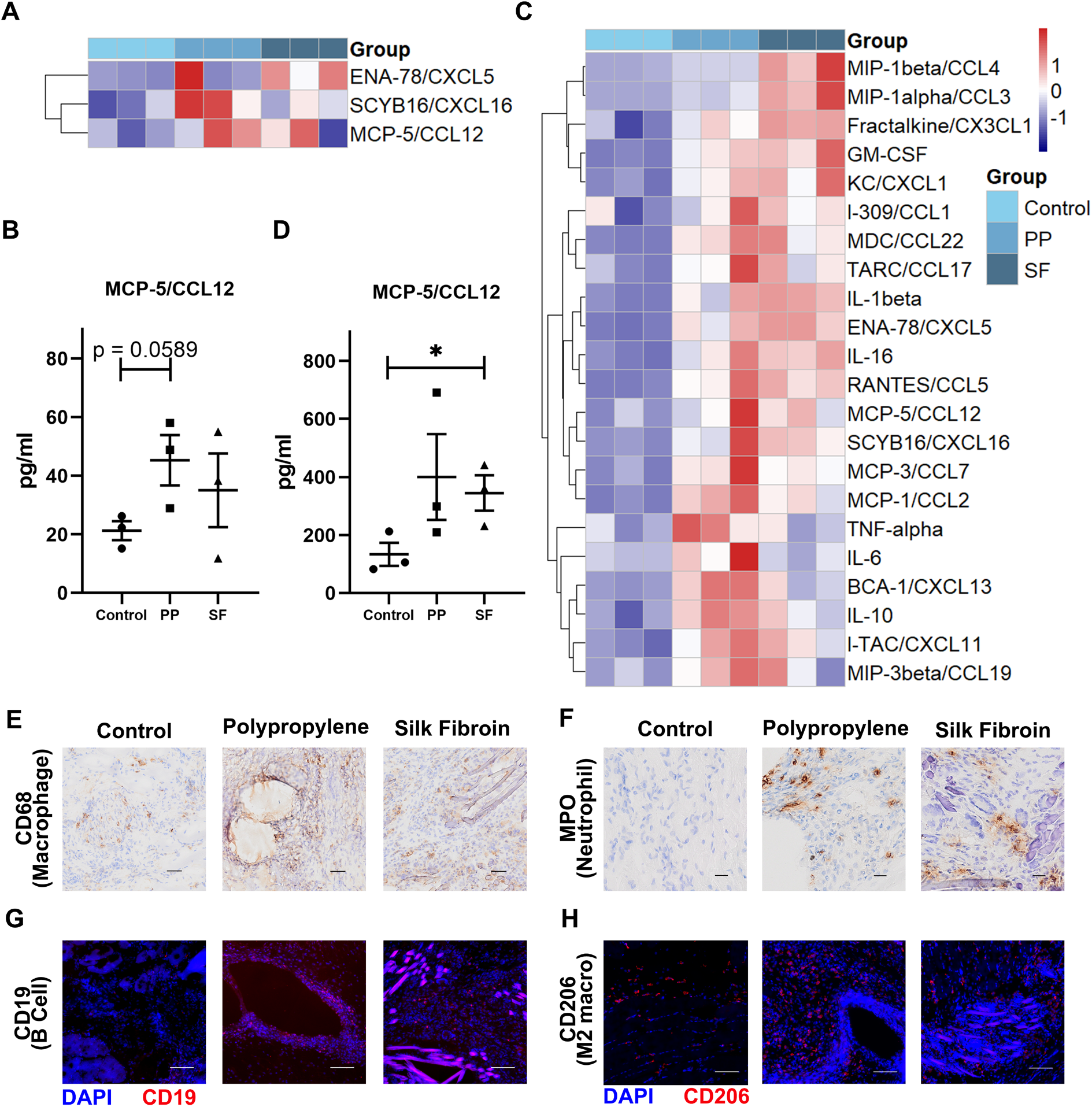
Inflammatory microenvironment surrounding the implant material caused an increase in blood circulating monocytes. **A**, Cytokine/chemokine concentrations were tested in the serum. **B**, MCP-5/Ccl12 chemokine concentration in the serum. Data are mean ± SEM (n = 3). Student’s t test. **C**, Cytokine/chemokine concentrations were tested in the muscle around biomaterials. **D**, MCP-5/Ccl12 chemokine concentration in the muscle around biomaterials. Data are mean ± SEM (n = 3). *p < 0.05, Student’s t test. **E-H**, Immunohistochemistry and immunofluorescence staining showing innate immune macrophage marker CD68, M2 macrophage marker CD206, Neutrophil marker MPO and adaptive immune B cells marker CD19 around biomaterials.

To identify the immune cells responsible for Ccl12 secretion, we performed immunohistochemistry and immunofluorescent staining surrounding the implant biomaterials. Specific markers for innate immune cell (macrophages, CD68, CD206; neutrophils, MPO) and adaptive immune cell (B cells, CD19) were identified. These results showed that both innate and adaptive immune cells increased surrounding mesh materials, as compared with controls (Fig. 4E-H), which become a potential candidate to activate blood circulation and systemically immune response.

In summary, biomaterials implantation aroused inflammatory microenvironment to initiate the increase of blood circulating monocytes, followed by recruiting into the liver and leading to lipid deposition.

### Subhead 5: Degradability of biomaterials decided local and remote organs response during a long-term implantation

The “foci” of the host remote organs responses is the inflammatory microenvironment surrounding biomaterials. In the short-term (2w and 6w), inert synthetic PP with nature-derived biological SF implantation showed the same implanted site and remote organs response pattern, while biodegradable biomaterials undergo continuous digestion and absorption in vivo. Thereby, we compared PP and SF in a situation of long-term (24w) implantation, and characterized their host responses locally and systemically.

As expected, SF showed continuous degradation while PP persist in situ with the assessment of HE (Hematoxylin-Eosin) staining (Fig. 5A, B). And from macroscopy, SF has degraded to invisible while PP remains (Supplementary Fig. 7A). Masson’s trichrome staining also suggested that SF showed decreased fibrosis while PP exhibited long-term fibrous encapsulation after 24w implantation (Fig. 5C). To assess the temporal patterns of inflammatory cell infiltration and immune responses in the implanted site, bulk RNA-seq and gene set enrichment analysis (GSEA) were performed. We summarized the normalized enrichment scores (NESs) for immune response and acute response related gene sets across all time points. Of note, the two groups display differences in 24w. Non-degradable synthetic PP preserved a high level of inflammation in 24w, while biological SF did not (Fig. 5D, red rectangle, inclined to blue), suggesting that SF shows long-term lower local host responses.

**Fig. 5.**
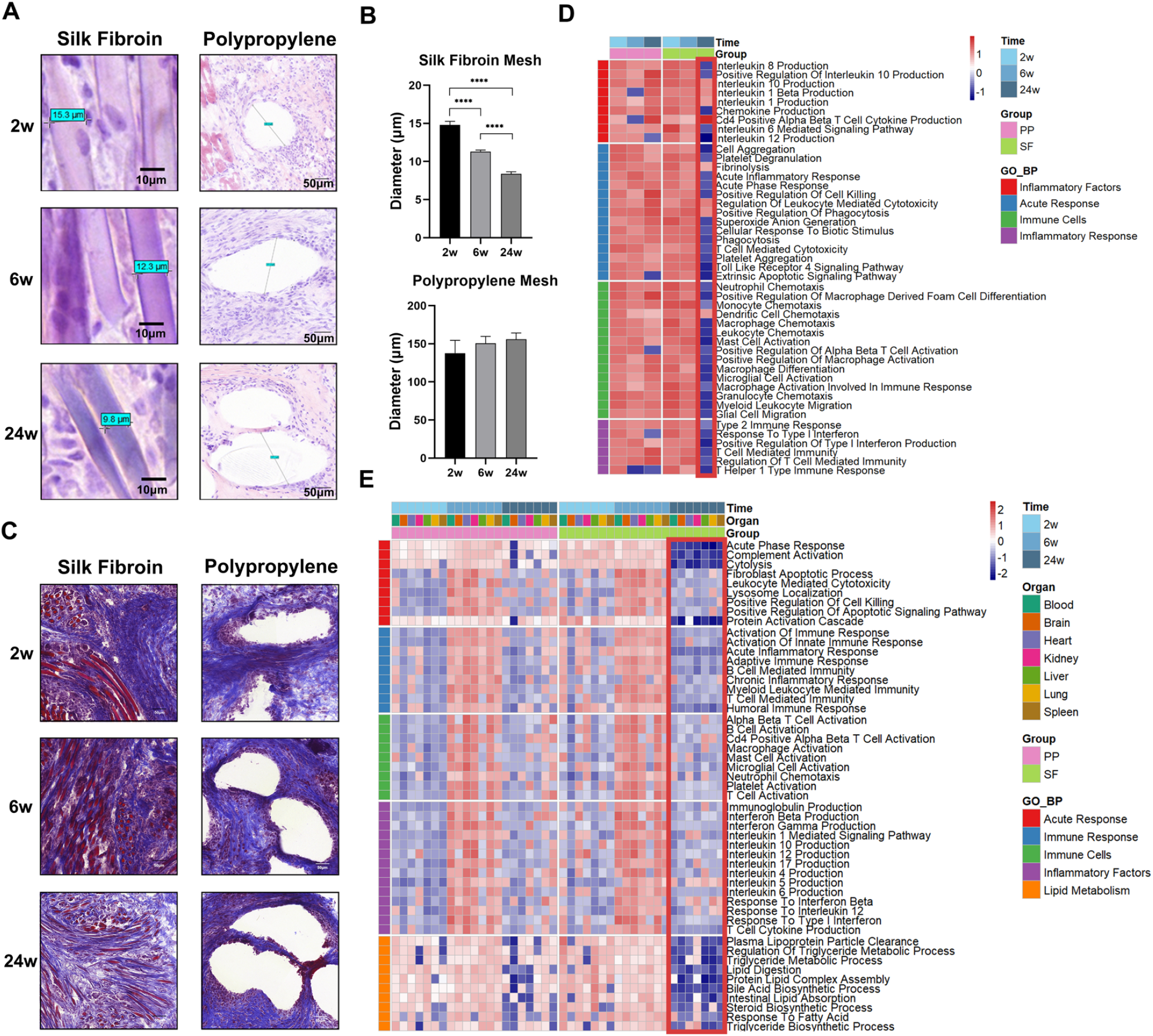
Degradable biological biomaterials can attenuate local and systemic response in the long-term. **A, B**, H&E staining and quantification of the materials diameter. Data are mean ± SEM (n = 14). ****p < 0.0001, Student’s t test. **C**, Masson’s staining of the tissue around biomaterials. **D**, Heatmap of NESs of tissue around biomaterials. **E**, Heatmap of NESs across all organs and time points.

For uncovering the temporal patterns of the remote organs responses, we performed a systematic assessment using GSEA across all organs and time points. We summarized the NESs for immune response and acute response related gene sets across all time points. We first counted the frequency of functional gene categories across all time points and organs, and found that the most commonly increased functional categories were related to acute response (coagulation, protein activation cascade and fibrinolysis), rejection response (xenobiotic metabolism and allograft rejection) and immune response (interferon gamma response and interferon alpha response) (Supplementary Fig. 7B, C; Supplementary Data 9-12), which were highly consistent with our previous analyses of DEGs clusters (Fig. 1D, E). We then plotted NESs for the immune response, acute response and lipid metabolism related GO biological process gene sets across all organs and time points. In both groups, immune response, acute response and lipid metabolism process occurred at 2w and enhanced at 6w (inclined to red) (Fig. 5E). Besides, immune cells were broadly consistent with inflammatory factors, reflecting that various immune cells activation are regulated by inflammatory factors (Fig. 5E). In 24w, an obvious decrease of selected NESs of SF group was observed (red rectangle, inclined to blue) (Fig. 5E). Notably, lipid metabolism related gene sets also showed the same temporal characteristics. Further fluorescence staining of liver were conducted at 24w and confirmed that lipid deposition was vanished in SF group and was aggravated in PP group after 24w implantation (Fig. 6A, B).

**Fig. 6.**
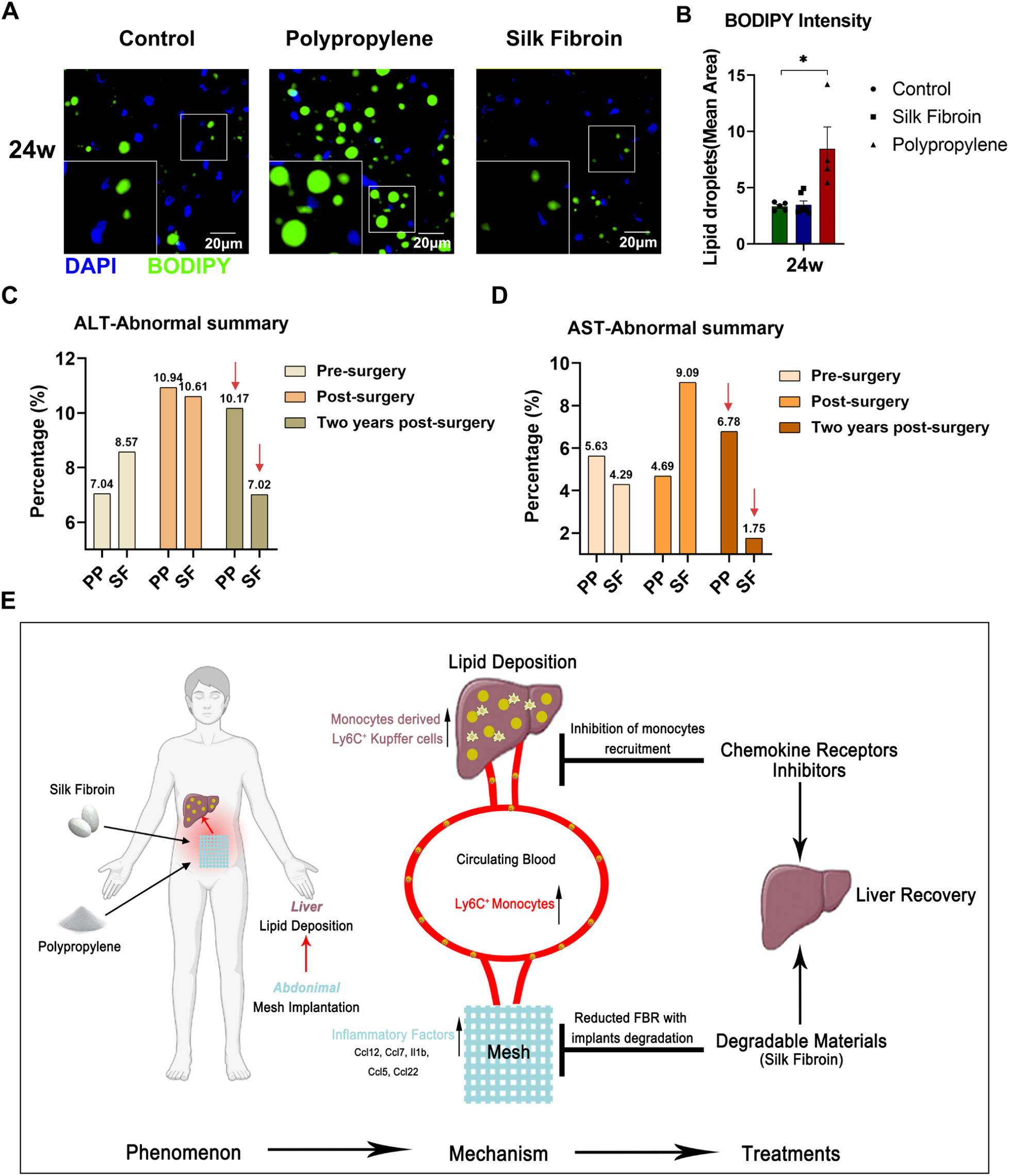
Clinical implantation of surgical meshes verified the time-specific hepatic lesion in patients. **A, B**, Fluorescence staining and quantification of the liver lipid after 24w implantation. Data are mean ± SEM (n = 4-7). *p < 0.05, Student’s t test. **C, D**, Frequency of abnormal ALT and AST value in all patients across 3 time points. **E**, Schematic overview of abdominal biomaterials implantation induced liver lipid deposition.

These results indicated that the disturbance of hepatic lipid metabolism persisted only in PP group after 24w implantation. Taken together, in comparison with non-degradable synthetic PP, degradable biological SF-based biomaterials offered the better capacity to minimize local and systemic response during long-term implantation.

### Subhead 6: The heterogeneity of hepatopathy in clinical implantation tightly associated with the degradability of biomaterials

To access whether patients with biomaterials implant show similar hepatic disorder, a randomized, double-blind, multicenter clinical trial were performed. 141 participants with abdominal hernia were enrolled and randomly assigned to the PP mesh (71 patients) and SF mesh (70 patients) groups. All patients received blood biochemical examinations pre-surgery, post-surgery (within one week) and two years post-surgery.

AST and ALT are strong indicators of hepatocyte function. We summarized the frequency of abnormal value (criteria: AST > 40U/L, ALT > 40U/L) in all patients across 3 time points. Compared with pre-surgery, the percentages of abnormal ALT elevated from 7.04% to 10.94% in PP group and from 8.57% to 10.61% in SF group post-surgery (Fig. 6C), implying that abdominal biomaterials implantation may also cause hepatic disorder in patients. Besides, SF group showed lower percentages of patients with abnormal ALT (10.17% versus 7.02%) and AST (6.78% versus 1.75%) than PP group after two years post-surgery (Fig. 6C, D). After a long-term implantation, degradable SF materials avoided the occurrence of hepatic disorder in patients (Fig. 6E).

## DISCUSSION

Taking the perspective of the whole organism, we described that remote major organ responds to local PP and SF mesh implantation for the first time. A priori, we expected remote organs response to local abdominal mesh implantation. A previous study demonstrated that the host evoked strong inflammatory activity in the remote organs such as the lung, spleen and small intestine after the subcutaneous implantation of gelatin-based conduit on the mice back*(25)*. In addition to that, our previously published study characterized two biomaterials-induced immune response in a rodent model in PBMC*(6)*, which indicated that the host interacts with implants systematically. Indeed, we found that all observed organs exhibited differential gene expression in at least one time point and immune responses were extensively activated, indicating that the immune response plays a predominant role in biomaterials-induced systematical damage. Moreover, these results emphasized the importance of systematical FBR assessment in new biomaterials development, which is still short in recent research. Our exploration documents the diversity of different organ responses to biomaterials implantation. We speculated some assumptions that can explain them. First, the baselines of gene expression profiles vary across organs*(34)*, affecting cellular reaction to biomaterials implantation stimuli. Second, the response of the organs may be affected by the interaction between immune cells and stromal cells, while the content of these cell types varies in different organs. Third, certain biomaterials implantation may have feature effects on specific organs. The comparison between natural-derived materials and synthetic materials has always been an important issue in the field of biomaterials. Some studies have shown that natural-derived materials are superior to synthetic materials in anti-infection*(35)*, promoting wound healing*(11, 17, 36)*. In this article, the duration of natural-derived and synthetic biomaterials induced remote organs responses were also investigated. Natural-derived SF mesh showed acute and immune responses in the short-term, but presented as safe and stable materials in the long-term. Synthetic PP is conventionally classified as an inert stable material; however, demonstrated sustained “active” in vivo. Clinically, surgical implantations are usually permanent*(37)*. The foreign implants induced discomfort in patients and complaints usually occurred years post-surgery, such as polyester mesh for hernia repair and silicone breast implants*(27, 38)*. Collectively, our results and clinical application scenarios suggested that long period host FBR assessment is necessary for biomaterials with clinical translation potential.

Of course, we are not emphasizing that the faster the biomaterials degradation, the better. All degradable materials should maintain their essential mechanical strength in the process of achieving tissue engineering function. Specifically, the degradable materials should gradually be replaced with the patients’ own cells and ECM over the course of recovery. As for SF-based materials, the rate of SF degradation is being managed by adjusting its molecular mass, crystalline size, or cross-linking*(39)*. In our previous study, SF-based scaffold was used to anterior cruciate ligament reconstruction and showed sustain mechanical support up to 18 months*(40)*. In this study, all hernia patients received SF mesh treatment showed satisfactory mechanical maintenance and low recurrence rate after two years (data not shown).

Nevertheless, there are some limitations in this study that need to be addressed. First, the types of biomaterials investigated are small. There is a huge number of materials available and biomaterials with different properties may display different systemic response patterns. Second, heart and lung also showed high DEG number and/or extensive pathway alteration, which further researches need to be carried out. There are several cells (macrophages, monocytes, and neutrophils, CD4+ T cells and CD8+ T cells) altered in blood after implantation*(6)*, which may mediate the heart and lung lesion. Third, a time point within two weeks was omitted. A shorter time point after implantation can help to understand the early stages of the response and how the innate immune response drives later tissue responses. Another open question is how local biomaterials implantation cause an increase in blood monocytes. Ccl12 may be the signal medium, however, further loss-of-function and gain-of-function experiments remain to be addressed.

## MATERIALS AND METHODS

### Study design

The primary goals of the study were to identify the systemic response profile after local biomaterials implantation and to provide a new reference for clinical biomaterials selection. Rats and mice were randomly assigned to all experiments. All animal studies were performed with approval by the Zhejiang University Institutional Animal Care and Use Committee (ZJU20220199). Whole-organism-wide RNA sequencing (RNA-seq) was performed on 7 major organs/tissues (blood, brain, heart, kidney, liver, lung, spleen) collected at 3 time points (2w, 6w, 24w) from rats with PP or SF mesh implantation and sham-operated controls. Each tissue in different group/time point was collected from the same region which outlined in Supplementary Fig. 8 and the principle of region selection is to avoid the influence of organ spatial heterogeneity on the results (try to choose a region that is homogeneous or contains all spatial structures). Downstream bioinformatic analysis was performed to identify organ-specific and temporal response.

### Surgical meshes

Polypropylene meshes (Prolene, Johnson-Johnson Inc., USA) were knitted from non-absorbable polypropylene monofilaments, which are resistant to enzymatic degradation and can maintain strength for clinical use. Silk fibroin meshes were purchased from Xingyue Co., Ltd. (Hangzhou, China), and the fabricate procedure was described in previous research*(24)* (Supplementary Fig. 7A(a)).

### Animal experiments

Rats and mice were fasting overnight before surgery. Animals were subcutaneously injected with pentobarbital (1mg/kg) and intramuscularly injected with Zoletil 50 (50mg/kg). After anesthesia, the abdominal hair was shaved and disinfected. Abdominal incision was performed, and the abdominal muscle layer was bluntly separated. Mesh (2×2 cm^2^ for rat and 0.75×0.75 cm^2^ for mice) was implanted between the peritoneal wall layer on one side of the abdominal wall and the muscle. The abdominal muscle layer was first sutured, followed by the abdominal skin. After the operation, the animals were quickly transferred back to the experimental cage.

### Clinical study design and participants

A total of 141 patients aged from 19 to 86, male or female, undergoing hernia repair were enrolled for this clinical trial, which conducted at the Xiangya Hospital of Central South University (Ethics Approved No. 201407070), Xiangtan Central Hospital (Ethics Approved No. 20140925), Xiangyang Central Hospital (Ethics Approved No. XYSZXYY-LL-PJ-2014-002), Huanggang Central Hospital (Ethics Approved No. HGYY-2014-010), the Fourth Hospital of Changsha (Ethics Approved No. CSSDSYY-LL-SC-2014-02) and the Second People’s Hospital of Yibin (Ethics Approved No. 2016-010-02) with all participants signed the subjects’ informed consent.

### Histological assessment

The specimens from 5 major organs/tissues (heart, kidney, liver, lung, spleen) were fixed by 4% paraformaldehyde and then paraffin embedded, and the muscle was harvested for cryotomy. Histological sections were prepared and subsequently subjected to hematoxylin and eosin (HE) staining and Masson trichrome staining.

### Liver lipid staining

Livers were collected, fixed in 4% paraformaldehyde and then dehydrated in sucrose solution (30% in PBS) at 4°C. Livers were then embedded with OCT (ThermoFisher) and sectioned. After hydration in PBS, frozen sections were incubated with Bodipy (0.1mg/mL in PBS) for 30 min to label lipids. After washing in PBS, the sections were counterstained for nuclei with DAPI (Beyotime).

### Blood routine examination

Intravenous blood samples from rats after surgery were collected in anticoagulative tube with EDTA-K2. Blood routine examination was carried out with fully automatic hematology analyzer XT-2000i (SYSEMX).

### Flow cytometry

Antibodies were purchased from BioLegend and Thermo Fisher Scientific. The following markers and clones were used: CD45 (Clone 104, 109814), CD11b (Clone M1/70, 101236), CD64 (Clone X54-5/7.1, 139311), Ly6G (Clone 1A8, 127623), Ly6C (Clone HK1.4, 128011), F4/80 (Clone BM8, 12-4801-82). Cell suspensions were stained with appropriate antibodies for 30 min on ice. Data were acquired on a BD LSRFortessa flow cytometer (BD Biosciences) and analyzed with FlowJo software (Tree Star). The gating strategies for the different populations are outlined in Supplementary Fig. 5.

### RNA-seq

RNA-seq was performed according to our previous method*(41)*. Briefly, RNA was extracted from samples by Trizol reagent (TAKARA), reverse transcription was conducted by SuperScript II reverse transcriptase (Invitrogen), double strand cDNA was synthesized using NEBNext mRNA second strand synthesis kit (NEB), double strand DNA was washed with AMPure XP beads (Beckman Coulter), and the sequencing library was constructed with Nextera XT kit (Illumina) and sequenced on Illumina X-Ten platform. RNA-seq reads data were mapped to a reference genome (Rnor_6.0(GCA_000001895.4)) from the Ensembl database using STAR*(42)*.

### Immunofluorescence

After the animals were sacrificed at 2w, 6w and 24w post-surgery, immunofluorescence was used to characterize the immune cell infiltration. The primary antibodies are listed as follows: CD68 (Bio-Rad, MCA341GA), CD206 (Abcam, ab64693), MPO (Abcam, ab208670) and CD19 (Abcam, ab245235). Stained histology sections were scanned under Olympus VS200.

### Magnetic luminex screening assay

Muscle samples or serum samples from the control and treatment groups were collected after 2w, and the concentration of cytokines and chemokines in each group was analyzed by magnetic luminex screening assay (R&D Systems).

### Protein-protein association networks

We submitted the DEGs names to the StringDB website (https://string-db.org/). Then, we obtained all interaction nodes with the lowest interaction score (0.4, the default setting). We removed less than four connected nodes and visualized them with Cytoscape (https://cytoscape.org/).

### Statistics

GraphPad Prism software was used for data analysis. Parametric data was analyzed by unpaired, two-tailed t test. DEGs was calculated by edgeR. For the overview in Figure 1B, C and Supplementary Data 1 - 4, differentially expressed transcripts (q-value < 0.1) were converted into unique gene symbols, and transcripts that did not map to the official name were removed. Figure 1D was generated with clusterProfiler. To functionally annotate clusters in Figure 1E, topGO was used. For GESA, RMA-normalized data was converted into GCT-file format and analyzed using javaGSEA Desktop Application (available from http://software.broadinstitute.org/gsea). We used gene set permutation with Signal2Noise as a ranking metric and the weighted scoring scheme. Other parameters were default.

## List of Supplementary Materials

Fig. S1 to S8 for multiple supplementary figures

Data file S1 to S12 (Excel files)

## Acknowledgments

We thank Zhang Tao, Lei Tingyun, Gu Yuqing, Zhang Zheyuan, Xie yuan and Li Jiajin for animal experiments and sample collection.

## Author contributions

Conceptualization: HWOY, XHZ, ZP, JJH, HYW

Methodology: ZP, CX, SCJ, JCY, BBW, HYW, RJL, YW, JHH

Investigation: ZP

Visualization: ZP, SCJ

Funding acquisition: HWOY, XHZ

Project administration: HWOY, XHZ, JJH

Supervision: HWOY, XHZ, JJH

Writing – original draft: ZP, XDY, XZZ, JXL

Writing – review & editing: HWOY, XHZ, ZP, XDY, JXL

## Data and materials availability

All data are available in the main text or the supplementary materials. Raw RNA-seq data will be uploaded to the public database when inquired by reviewers or after publication.

