## Supplementary Figures for "Biomaterial based implants caused remote liver fatty deposition through activated blood-derived Kupffer cells"

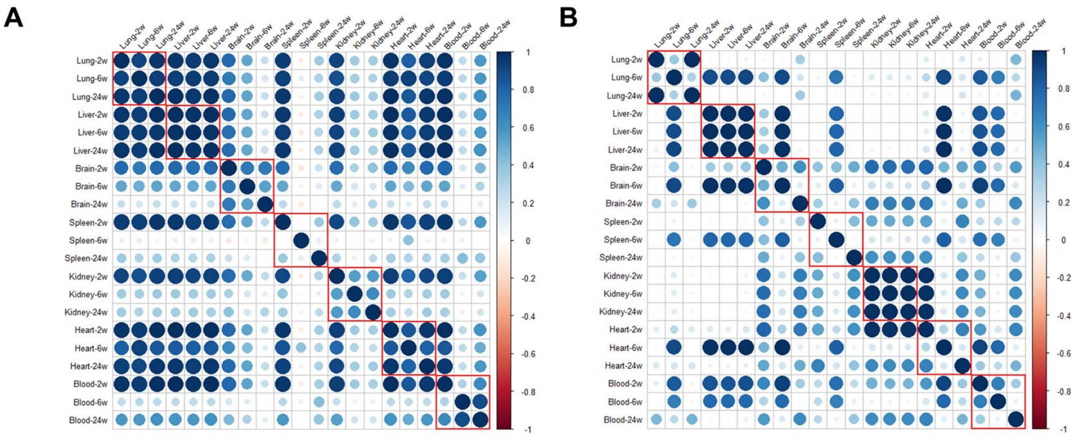


**Supplemental Fig.1 | Similarity of differentially expressed gene within multi-organs.** Pairwise comparison between genes with fold change >2 in PP (**A**) group and SF (**B**) group.

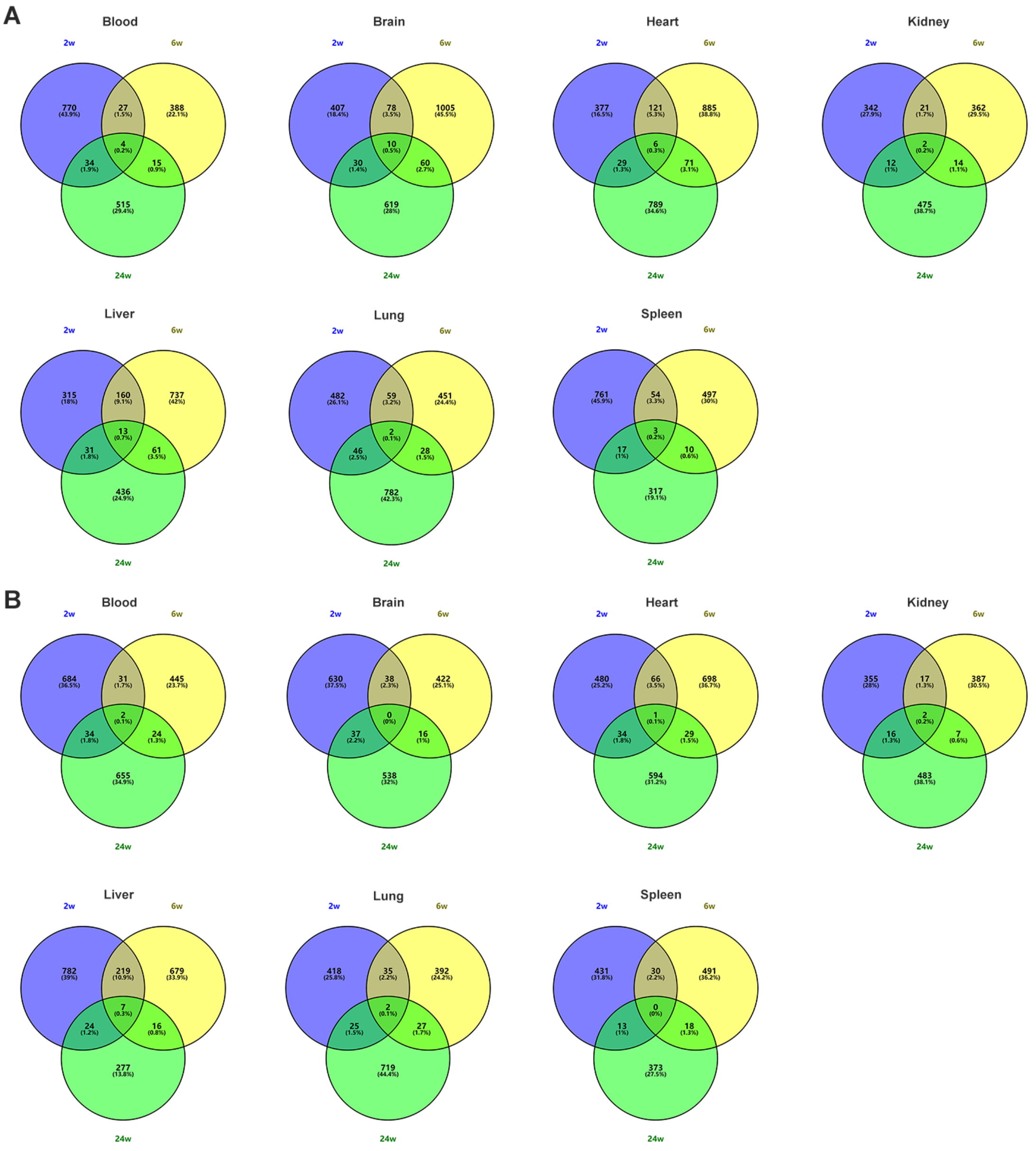


**Supplemental Fig.2 | All genes that had a fold change >2 in at least two timepoints.** Venn diagrams of genes with fold change >2 in PP (**A**) group and SF (**B**) group.

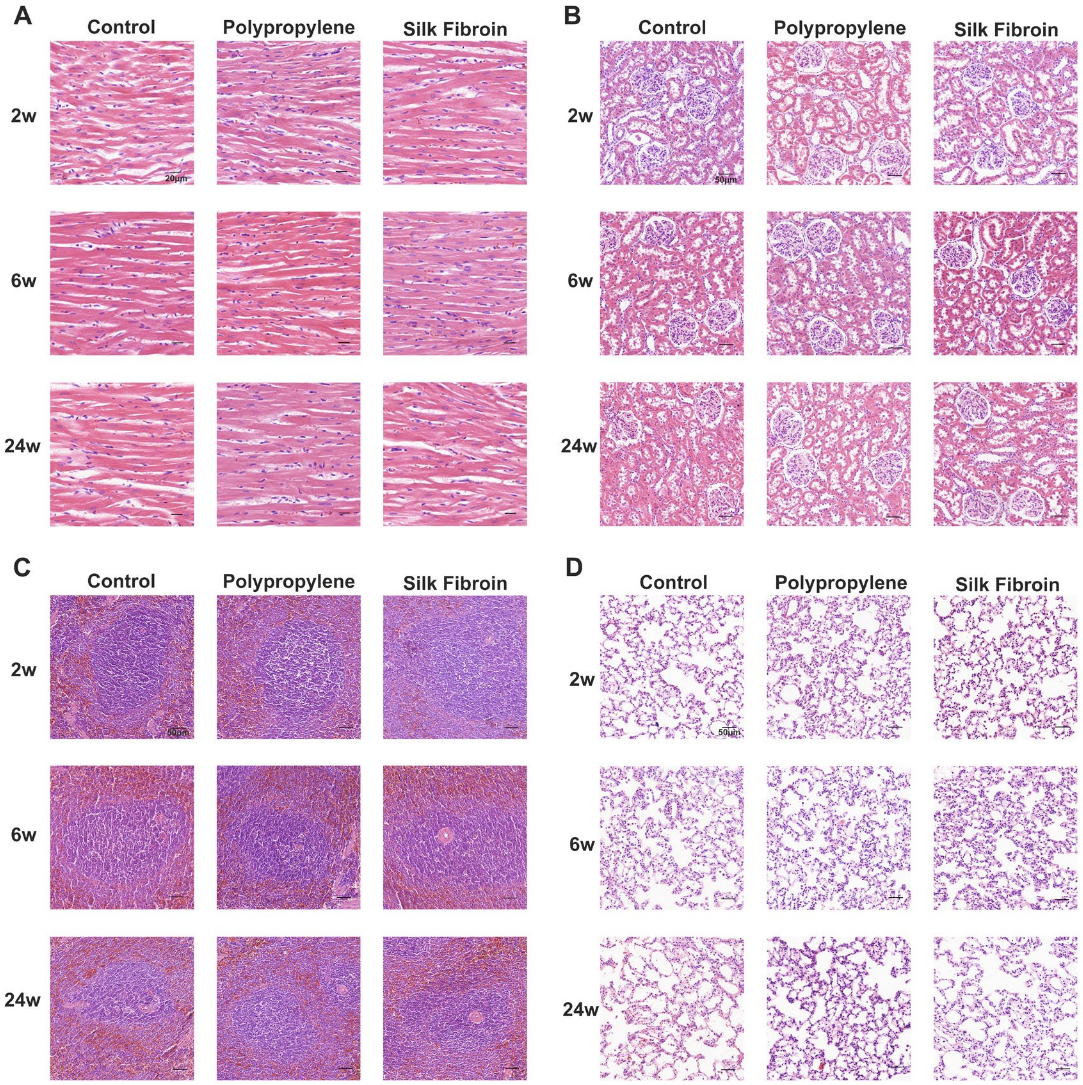


**Supplemental Fig.3 | No abnormalities were found with histological morphology evaluation of major organs.** Hematoxylin-Eosin staining of heart (**A**), kidney (**B**), spleen (**C**) and lung (**D**).

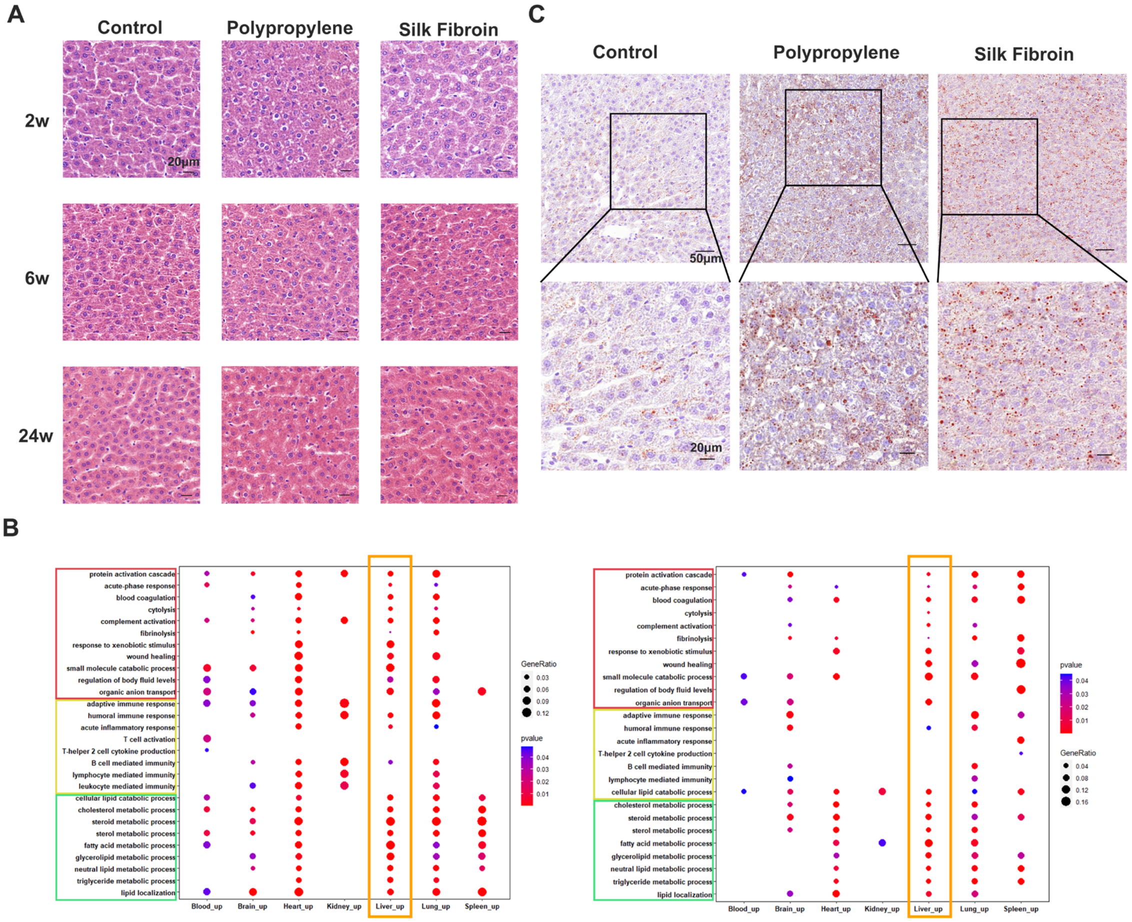


**Supplemental Fig.4 | Liver function pathways were extensively interfered and showed lipid droplets deposition.** **A**, Hematoxylin-Eosin staining of liver. **B**, GO enrichment analysis of genes that are expressed with a fold change > 2 in at least two time points. PP (left) and SF (right). **C**, Oil red O staining of liver sections after 2w mesh implantation.

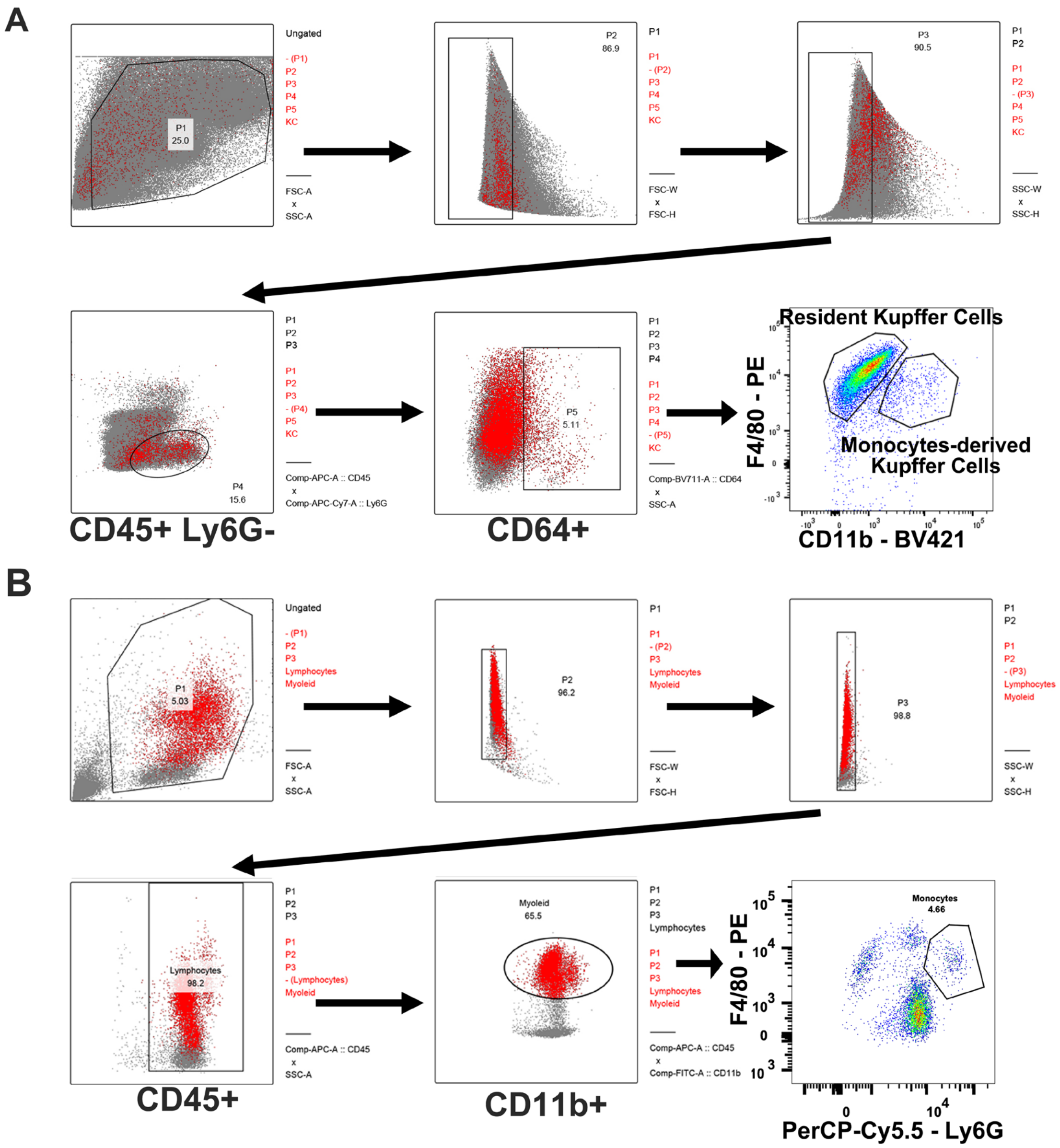


**Supplemental Fig.5 | Flow cytometry gating strategy for liver and blood. A**, Gating for resident Kupffer cells and monocyte-derived KCs in liver. **B**, Gating for monocytes in blood.

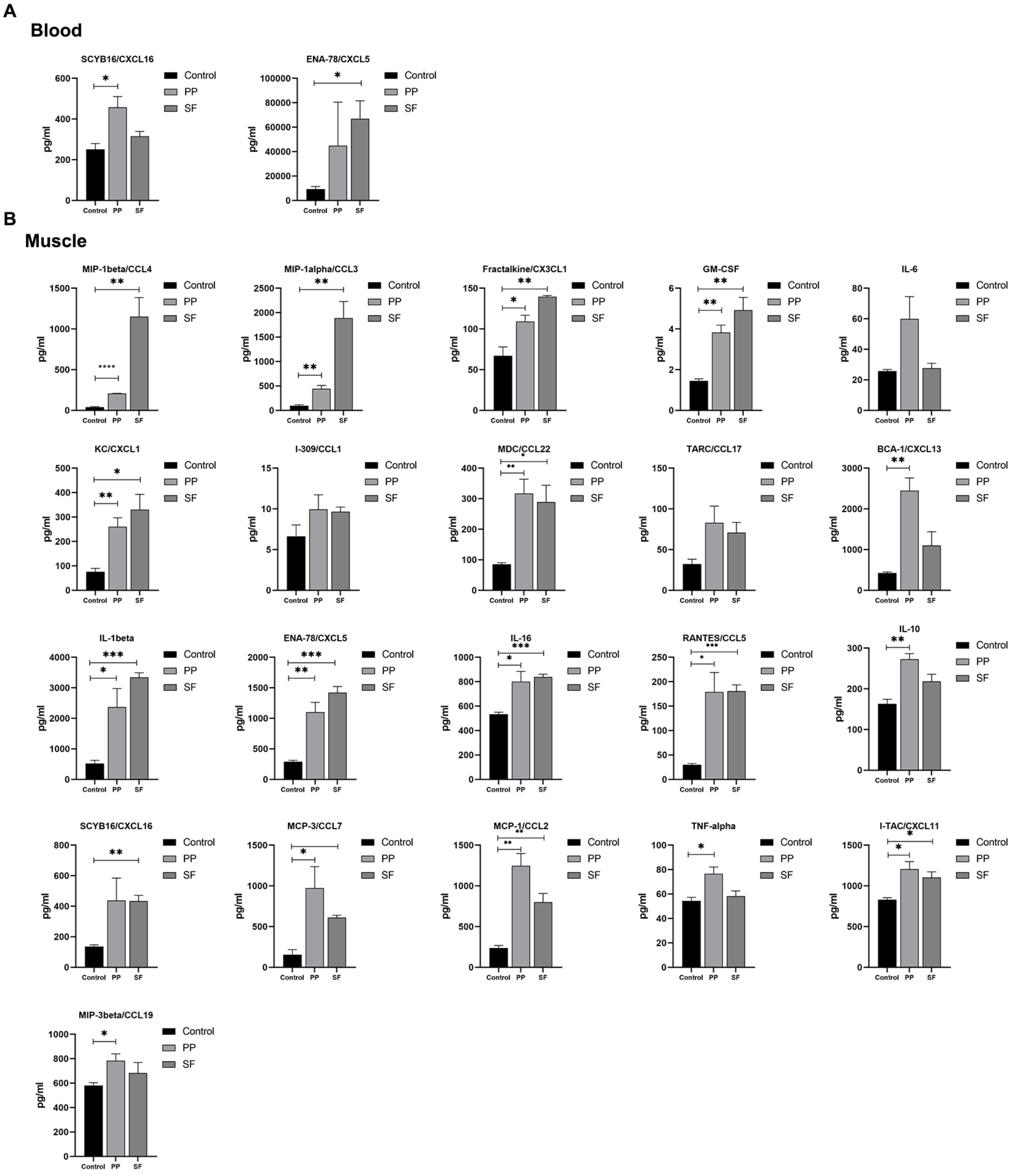


**Supplemental Fig.6 | The increase of inflammatory factors around the biomaterials promote the increase of monocytes in the blood.** Cytokines and chemokines levels in the blood serum (**A**) and surrounding the implant biomaterials (**B**). Data are mean ± SEM (n = 3). *p < 0.05, **p < 0.05, ***p < 0.05, Student's t test.

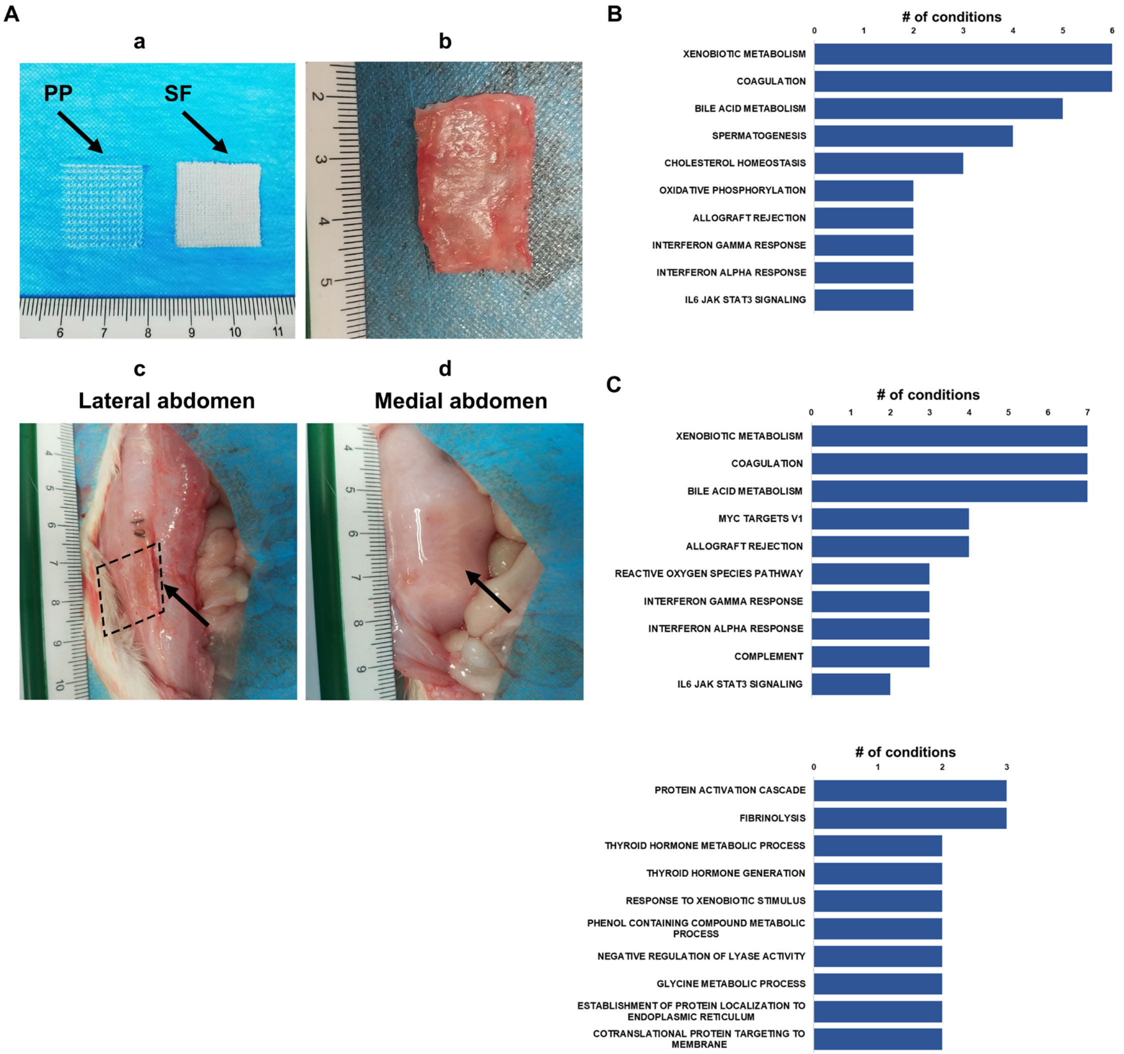


**Supplemental Fig.7 | PP mesh persist in situ after long-term implantation and the most commonly increased functional categories. A**, Macroscopy of implanted mesh after 24w. Frequency of functional gene categories across all time points and organs in PP (**B**) group and SF (**C**) group.

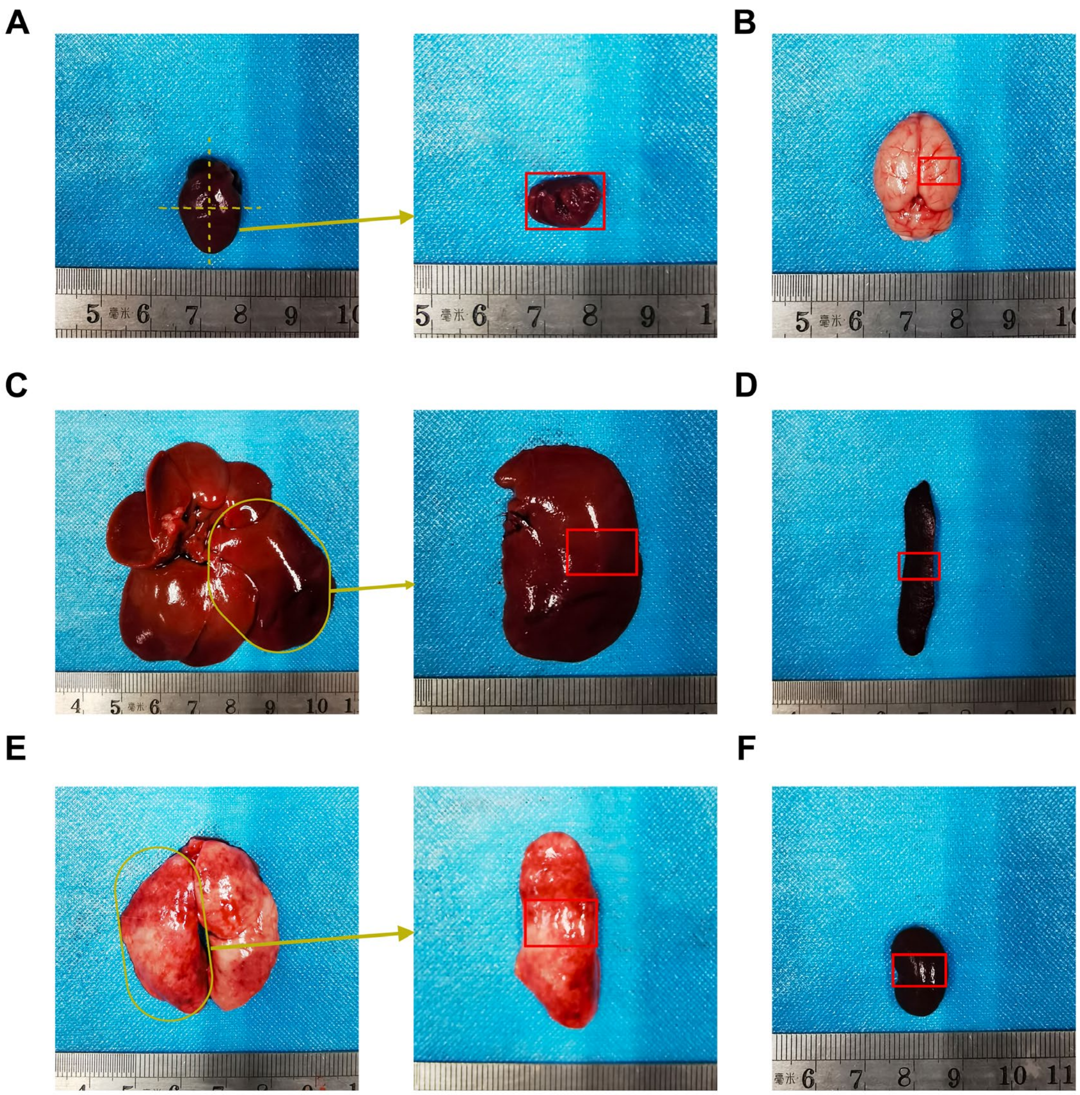


**Supplemental Fig.8 | Region selection of all investigated organs (red rectangle). A**, Heart collected the structure near the left ventricle. **B**, Brain collected the right cerebral cortex. **C**, Liver collected the middle area of the largest hepatic lobe. **D**, Spleen collected the middle area as red rectangle indicated. **E**, Lung collected the middle area of the largest lobe. **F**, Kidney collected the middle area as red rectangle indicated.
